# Acetaldehyde makes a distinct mutation signature in single-stranded DNA

**DOI:** 10.1101/2022.03.22.485364

**Authors:** Sriram Vijayraghavan, Latarsha Porcher, Piotr A Mieczkowski, Natalie Saini

## Abstract

Acetaldehyde (AA), a by-product of ethanol metabolism, is acutely toxic due to its ability to react with various biological molecules including DNA and proteins, which can greatly impede key processes such as replication and transcription and lead to DNA damage. As such AA is classified as a group 1 carcinogen by the International Agency for Research on Cancer (IARC). Previous *in vitro* studies have shown that AA generates bulky adducts on DNA, with signature guanine-centered (GG→TT) mutations. However, due to its weak mutagenicity, short chemical half-life, and the absence of powerful genetic assays, there is considerable variability in reporting the genotoxic effects of AA *in vivo*. Here, we used an established yeast genetic reporter system and demonstrate that AA is highly mutagenic and makes strand-biased mutations on guanines (G→T) at a high frequency on single stranded DNA (ssDNA). We further demonstrate that AA-derived mutations occur through lesion bypass on ssDNA by the translesion polymerase Polζ. Finally, we describe a unique mutation signature for AA, which we then identify in several whole-genome and -exome sequenced cancers, particularly those associated with alcohol consumption.

## Introduction

Alcohol consumption is associated with a variety of cancers and is among the leading causes of mortality in humans. One of the key metabolites driving alcohol toxicity is acetaldehyde (AA), which is generated by the reduction of ethanol. Additionally, AA can be obtained from a variety of other foods and beverages, as well as tobacco smoke (1). *In vivo*, AA is oxidized to acetate through a series of NAD+-dependent aldehyde dehydrogenases ((2) and reviewed in (3)). However, dysfunction of AA-detoxifying mechanisms has severe cytotoxic and mutagenic outcomes, which can lead to carcinogenesis. Free AA is highly reactive towards key biomolecules including DNA and protein, which can inhibit cellular processes and contribute to carcinogenesis (4,5). Based on its toxic properties, the International Agency on Research on Cancer classifies alcohol consumption-associated AA as a group I carcinogen (6).

The genotoxic effects of AA have been explored in several studies. Previous *in vitro* studies have demonstrated that AA can react with guanine residues on DNA and form bulky adducts (reviewed in (7)). Examples include the well-studied adduct N^2^-ethyl-2’-deoxyguanosine (N^2^-Et-dG) (8), which has been often used in *in vitro* studies to analyze the effects of AA-mediated DNA damage (9–11). In human embryonic kidney cells, N^2^-Et-dG are associated with GC→TA transversions (12). N^2^-Et-dG has been shown to stall replication (12), as well as transcription (13) Additionally, other AA-derived DNA adducts such as α-S- and α-R-methyl-γ-hydroxy-1, N2-propano-2’-deoxyguanosine (CrPdG) have been shown to induce DNA intra-strand crosslinks in adjacent guanine residues, leading to signature GG→TT transversions in single-and double-stranded plasmid DNA, as well as human fibroblast cell lines (14). Acetaldehyde has been shown to induce chromosomal-scale DNA damage in CHO cells deficient in homologous recombination and nucleotide excision repair (15). Studies in both budding yeast and fission yeast have genetically demonstrated the induction of DNA repair pathways by AA treatment (16,17). Using *Xenopus* egg extracts, it was shown that AA-derived DNA inter-strand crosslinks retard replication fork progression and lead to increased mutation frequency (18). In colorectal cancer cell lines, elimination of tumor-suppressor HR genes results in AA hypersensitivity (19), further highlighting AA-induced DNA damage and the ensuing cellular responses that sense such damage.

However, a diagnostic and specific mutagenic signature of AA exposure *in vivo* has proven elusive. Mutation signatures, which are the characteristic patterns of single and double base substitutions associated with discrete mutagens and metabolic processes, help understand the mutagenesis mechanisms leading up to cancer development. Advances in computational analyses of large-scale sequencing data has revealed the patterns of somatic mutations for thousands of whole-genome and- exome sequenced cancers by several groups (20–23). Therefore, identification of a discrete AA-associated mutation signature would serve as a critical predictor of carcinogenesis especially in alcohol-associated cancers. Previous studies investigating ethanol mutagenicity in yeast were similarly inconclusive as to the role of AA in mediating mutagenesis as AA was not found to be mutagenic in these yeast strains (24). Treatment of induced pluripotent stem cells (IPSCs) with AA led to a profound DNA damage response, however, no increase in mutagenesis was seen in these cells (25). In esophageal carcinoma patients, an alcohol-associated T→C mutation signatures has been described; although the mutation signature correlated with smoking and drinking, the signature was not specific for AA-induced mutations (26). Overall, while AA is demonstrably mutagenic, there is a lack of consensus among different studies as to its precise mutation signature and spectrum *in vivo.* An obvious commonality that could reconcile the above studies is that most *in vivo* analyses of AA mutagenicity is conducted in DNA-repair proficient backgrounds. As a result, low mutagenicity combined with efficient repair could easily confound an AA-specific mutation signature. Discrepancies between the induction of a DNA damage response, increased genome-instability and carcinogenesis associated with AA and the lack of detectable mutagenesis by AA necessitates sensitive models to unambiguously detect the mutations induced by AA.

In the present study, we investigate the mutation spectrum and signature of acetaldehyde using a sensitive yeast reporter assay. We show that AA strongly mutagenizes single-stranded DNA *in vivo* and can induce strand biased mutations at an elevated frequency. AA-derived mutations depend on translesion synthesis by DNA polymerase zeta (Polζ). Using whole-genome sequencing, we determined the mutation spectrum of AA and describe its trinucleotide mutation signature. Finally, we identified cancers among the whole-genome-sequenced cohorts from the Pan-cancer Analysis of Whole Genomes (PCAWG) (23) and whole-exome-sequenced cohorts from the International Cancer Genome Consortium (ICGC) (27) carrying the novel AA mutation signature and show that it positively correlates with a previously identified AA-associated mutation signature. Our work provides a novel and diagnostic signature of acetaldehyde exposure in yeast and cancers.

## Materials and Methods

### Yeast strains

Strains used in this study are derivatives of CG379 with the genotype *MATα his7-2 leu2-3,112 trp1-289, cdc13-1.* The genes *CAN1, URA3, ADE2* and *LYS2* were deleted from their original loci and reintroduced as the array *lys2::ADE2-URA3-CAN1* at the left de novo telomere arm of Chromosome 5, as described earlier (28). *RAD1, REV3* and *RAD30* were deleted using *KANMX.* The strain carrying the *rev1-AA* allele was the same as described earlier (29). All the strains and the primers used in the study are listed in Table S1.

### Acetaldehyde-induced mutagenesis

Yeast strains carrying the *cdc13-1* temperature sensitive (ts) allele were grown as described earlier (28). Briefly, cultures of the *cdc13-1* strains were grown at 23 °C for 72 hours until saturation. Roughly 10^7^ cells were inoculated into fresh YPD and grown with shaking at 37 °C for 6 hours in Erlenmeyer flasks to induce resection at telomeres. Cells were monitored for G2 arrest by microscopy, at which point >95% cells arrested as large double buds. Thereafter, cells were harvested by centrifugation, washed three times with sterile water and resuspended in water in 15ml conical tubes. Acetaldehyde was added to samples at a final concentration of 0.2%, and samples were incubated alongside the control samples (without AA) at 37 °C in a rotary shaker for 24 hours. To minimize AA evaporation during the experiment, 1) AA was kept on ice prior to addition, 2) pre-chilled pipette tips were used to dispense AA, 3) tubes will filled to the top, leaving very little headspace during rotary incubation, and 4) tubes were sealed with parafilm. Appropriate dilutions of cells were plated on complete synthetic complete (SC) media (MP Biomedicals) to measure viability and SC-Arginine plates containing 60mg/ml canavanine (Sigma) and 20mg/ml adenine to isolate Can^R^Ade^-^ mutants (red colonies). All plates were incubated at 23 °C for 5-7 days until colonies appeared. Mutants were tested for Chromosome V left arm loss by replica plating cells on SC-Uracil to select for loss of the *URA3* gene as described earlier (28). Genomic DNA was isolated from independent Can^R^Ade^-^ mutants for whole genome sequencing.

### DNA sequencing

Genomic DNA was isolated from yeast strains using the Zymo YeastStar genomic DNA isolation kit (Genesee Scientific) using the manufacturer’s recommended protocol. DNA concentrations were measured via Qubit (Invitrogen) and diluted to approximately 10ng/ul for library preparation. Diluted DNA was used for fragmentation on Covaris LE 220 system. The KAPA Hyper prep system (Roche) for library preparation with each sample acquiring a unique dual index adapter. After pooling all instances, the Illumina NovaSeq sequencing system for analysis. Sequencing reads were aligned to the reference genome ySR127 (30) using BWA-mem (31) and duplicate reads were removed using Picard tools (http://broadinstitute.github.io/picard/). Single nucleotide variants (SNVs) were identified using VarScan2 (32), using a variant allele frequency filter of 90%. Unique SNVs were by identified by comparing AA-treated samples with untreated parent strains serving as matched normal and removing duplicates.

### Mutation spectrum and signature analysis

Mutation analysis was done as previously described (29). Mutations within 30 kb of the chromosome ends were classified as sub-telomeric and those lying beyond were binned as mid-chromosomal. Mutation spectra was plotted as pyrimidine changes, taking into consideration reverse complements for each base change. SomaticSignatures (33) was used to plot mutation spectra across 96 channels to account for all possible base substitutions. Strand-biased mutations were evaluated based on whether the SNVs were located on ssDNA generated upon telomere uncapping and resection. Mutations per isolate were calculated by plotting SNV as a function of the total number of strains used per treatment condition. pLogo (https://plogo.uconn.edu/) was used to infer the mutation signature of AA treatment, by evaluating the statistical probability of over-/under-representation of residues in the ±1 trinucleotide context of the mutated residue compared to the background sequence. For a particular substitution, reverse complements were taken into consideration to perform the pLogo calculation.

### Mutation enrichment and mutation load analysis

Mutation enrichment and mutation loads were calculated based on (28,29) using Trinucleotide Mutation Signatures (TriMS) whereby the number of a given substitution in a specific trinucleotide context is compared against the total number of the given substitution genome-wide, as well the incidence of the mutated residue within the ±20 nucleotide context of the mutation. The following calculation was used:

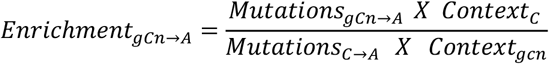

A one-sided Fisher’s Exact test was used to calculate the p-values of enrichment of the given mutation signature in each sample and in the total yeast samples. Mutation loads for a given signature were calculated with a minimum enrichment probability of >1 and a Bonferroni corrected p-value of ≥0.05, using the following equation:

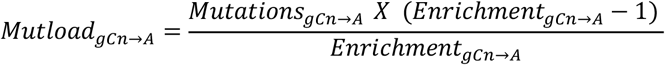

### Mutational analysis in cancers

Somatic mutation load and enrichment were calculated for a given signature using mutation data from deduplicated somatic SNV calls from different donors in whole-genome-sequenced cancers from PCAWG (23) and whole-exome-sequenced cancers from ICGC data portal (27). For cancer samples carrying an enrichment of the AA mutation signature of ≥1 (Bonferroni-corrected p-value of ≤0.05), transcriptional strand bias of mutations was calculated with BEDTools (34) intersect, using hg19 as the reference genome (UCSC Table Browser, (35)) and a goodness of fit test was performed to test the statistical significance of the ratios of mutations on transcribed vs non-transcribed strands. Mutation signature correlations were performed using simple linear regression and significance of correlations were estimated based on Pearson’s r coefficient.

## Results

### Acetaldehyde treatment results in elevated mutation frequency in yeast cells

#### A primer on the yeast mutation reporter system

Single stranded DNA (ssDNA) can be exposed during several key steps of DNA metabolism including DNA replication and transcription, and is highly susceptible to mutagens, which makes it an ideal template to uncover mutation profiles of even weakly mutagenic agents. To this end, we used a previously developed genetic reporter assay using the budding yeast *Saccharomyces cerevisiae*, whereby the strain has a temperature-sensitive (ts) *cdc13-1* allele, and has additionally been engineered to carry the *ADE2, CAN1* and *URA3* genes near the *de novo* telomere of the left arm of Chromosome 5 after being removed from their original chromosomal loci (Fig 1A, (28)). At the non-permissive temperature (37 °C), the *cdc13-1* allele causes telomere uncapping, resulting in extensive resection at chromosome ends and the production of large tracts of ssDNA (36). These tracts extend up to 30 kb from telomere ends and span the above reporter genes placed within the sub-telomeric regions (37,38). Additionally, induction of the ts allele causes yeast to arrest in G2 as large buds (39). Treatment of cells at this stage with a mutagen permits accrual of lesions within the exposed ssDNA and shifting of the strains to the permissive temperature (23 °C) hereafter results in resynthesis of the second strand with the mutagenic bypass of the lesions and produce selectable mutations (28). Due to the single-stranded nature of DNA, excision repair pathways cannot remove the lesions allowing us to analyze the signature of lesion formation in ssDNA. Mutations within the *CAN1* and *ADE2* reporter genes are selectable. *CAN1* mutations render yeast cells resistant to the arginine mimicking chemical canavanine (Can^R^), whereas mutations in *ADE2* fail to synthesize adenine and when placed on low adenine media, appear as red or reddish-pink colonies (Ade^-^). Clustering of mutations within these reporters produce Can^R^Ade^-^ mutants which are visually quantified to calculate the mutation frequency associated with the given mutagen. This system has been successfully employed to test the mutagenicity of multiple agents, including the APOBEC family of cytidine deaminases (28,40), and alkylating agents methyl-methanosulfonate (MMS) and ethyl-methanosulfonate (EMS) (29,41).

**Fig 1:**
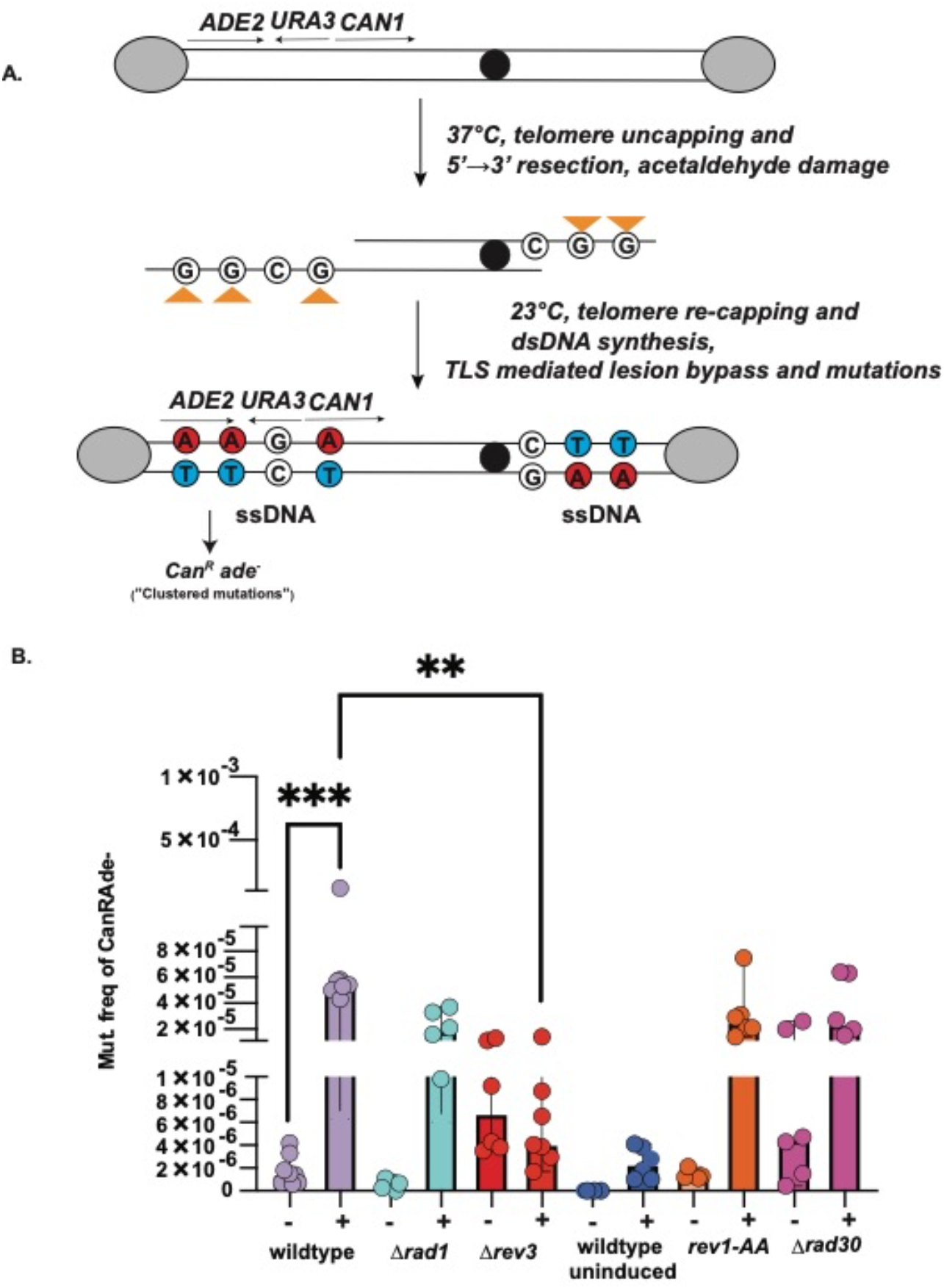
Testing the mutagenicity of acetaldehyde in yeast. A) Schematic of the yeast reporter system. As described previously (28), the assay utilizes a yeast strain harboring the *cdc13-1* allele and sub-telomeric reporters *CAN1, ADE2* and *URA3* on chromosome V (Materials and Methods). Temperature shift (37 °C) followed by AA addition produces lesions (orange triangles) on guanine residues within ssDNA within reporter genes, which upon restoration of permissive temperature (23 °C) would be erroneously bypassed by translesion synthesis, resulting in “clustered” Can^R^Ade^-^ mutations within the reporters (solid red circles). Grey ovals-telomere protection complex, solid black circle-centromere. B) Acetaldehyde mutation frequency estimates. Frequency of Can^R^Ade^-^ isolates after 24h AA exposure in the indicated strains. Minus (-) indicates water-treated controls whereas plus (+) indicates AA treatment. The “wildtype uninduced” dataset indicates mutation frequencies in the wildtype strain in the absence of *cdc13-1* induction (37 °C), with the entire assay conducted at ambient temperature (23 °C). Data represents median frequencies with 95% CI. Asterisks represent p-value <0.05 based on an unpaired t-test (wt vs *Δrev3* = 0.0005). Plotting and analysis were performed using Prism (v 9.3.1, GraphPad Software, LLC)

#### Acetaldehyde is mutagenic on ssDNA in yeast

We used the above reporter system to test if AA can induce mutations in yeast. To test this, upon induction of *cdc13-1* via shifting yeast cells to non-permissible temperature (37 °C), we treated G2-arrested yeast cells with AA. AA is extremely volatile, with a melting temperature of 18 °C. To minimize loss due to evaporation, we added chilled AA to cultures and additionally filled culture tubes to capacity to minimize AA oxidation during incubation. The strains were incubated with AA in a rotating incubator to allow the yeast cells to stay in suspension in the media and to allow maximum exposure to AA in the media. We did not notice any appreciable reduction in the viability of cells treated with 0.2% AA, compared to water-only control samples (Fig S1, Table S2).

AA treatment for 24 h led to a > 30-fold increase in mutation frequency (median mutation frequency 5.30E-05) over strains treated with water (median frequency 1.40E-06), Fig 1B). Since AA is highly volatile, we also determined the mutation frequencies of yeast treated with AA at 4 °C for 24h. AA was found to be equally mutagenic at this temperature as 37 °C as such, all further experiments were conducted at 37 °C (Table S2).

### Acetaldehyde mutagenesis relies on translesion synthesis

Since cells were treated with AA after the induction of the formation of ssDNA, the mutations were likely due to bypass of lesions during the resynthesis of the second strand to restore DNA to its doublestranded form. Such mutations should be independent of DNA repair by nucleotide excision repair (NER), which requires a homologous template (42). We deleted *RAD1* to abolish NER in strains and saw that there was no statistically significant change in the mutation frequency upon treatment with AA (Median mutation frequency 1.85E-05) (Fig 1B, Table S2), indicating that NER does not function to repair lesions in ssDNA.

In yeast the two major TLS pathways rely upon either DNA polymerase eta (Pol η) which is involved in error-free bypass of lesions, or the error prone DNA polymerase zeta (Pol ζ). The latter additionally involves Rev1, which in addition to being a structural component of Pol ζ can independently catalyze bypass of certain types of lesions (43–45). To test which of the above pathways contributes to the elevated mutation frequency observed with AA, we deleted *RAD30* which codes for the catalytic subunit of the Pol η polymerase. There was no reduction in mutation frequency from AA treatment in *Δrad30* strains compared to the wildtype strains (Median mutation frequency 2.35E-05) (Fig. 1B, Table S2). Next, we deleted *REV3* in our strains, which encodes the catalytic subunit of Pol ζ. In the *Δrev3* background, AA treatment caused a 13-fold reduction in mutation frequencies compared to the wildtype strains (Fig. 1B, Table S2). Lastly, we assessed the effect of Rev1 (29). The catalytically-inactive *rev1-AA* mutant (D647A and E648A, (44) did not alter the mutation frequency with AA treatment (Fig 1B, Table S2). Our results strongly suggest that erroneous bypass of lesions in ssDNA by Polζ underlie the mutagenicity of AA.

### Acetaldehyde predominantly makes G→T transversions on ssDNA

We next asked if we can identify the mutation spectrum associated with exposure to acetaldehyde. For this, we isolated genomic DNA from 115 Can^R^Ade^-^ mutant colonies obtained from AA treatment and performed whole genome sequencing. AA treatment leads to the formation of a lesion on ssDNA whose erroneous bypass further leads to mutations culminating as Can^R^Ade^-^ isolates. However, Can^R^Ade^-^ isolates can also arise from the loss of the chromosomal arm carrying the reporters. To rule out this possibility, we additionally tested the Can^R^Ade^-^ isolates derived from AA treatment for the presence of the *URA3* gene within the reporter and only selected Can^R^Ade^-^ *Ura^+^* isolates in the AA treatment cohort for further analysis (Table S3).

To identify mutations specific to AA treatment, we sequenced and analyzed 27 isolates derived from control (water) treatment in parallel. Compared to mutagen treated samples, control treatments do not result in the appearance of a high frequency of red Can^R^Ade^-^ (red) colonies, therefore we sequenced a randomized mixture of Can^R^ (white) and Can^R^Ade^-^ (red) colonies (Table S2). Unlike samples treated with water, among the 504 unique mutations identified in AA-treated samples, C→A (G→T) transversions were the predominant SNVs (41%), along with a smaller percentage of C→T (G→A) changes (6.7%) (mutation density for cumulative C→A changes per isolate, water=0.59, AA=1.79), Fig 2, Table S4). Our observations are in accordance with prior studies showing guanines to be the primary target of AA-induced lesions.

**Fig 2:**
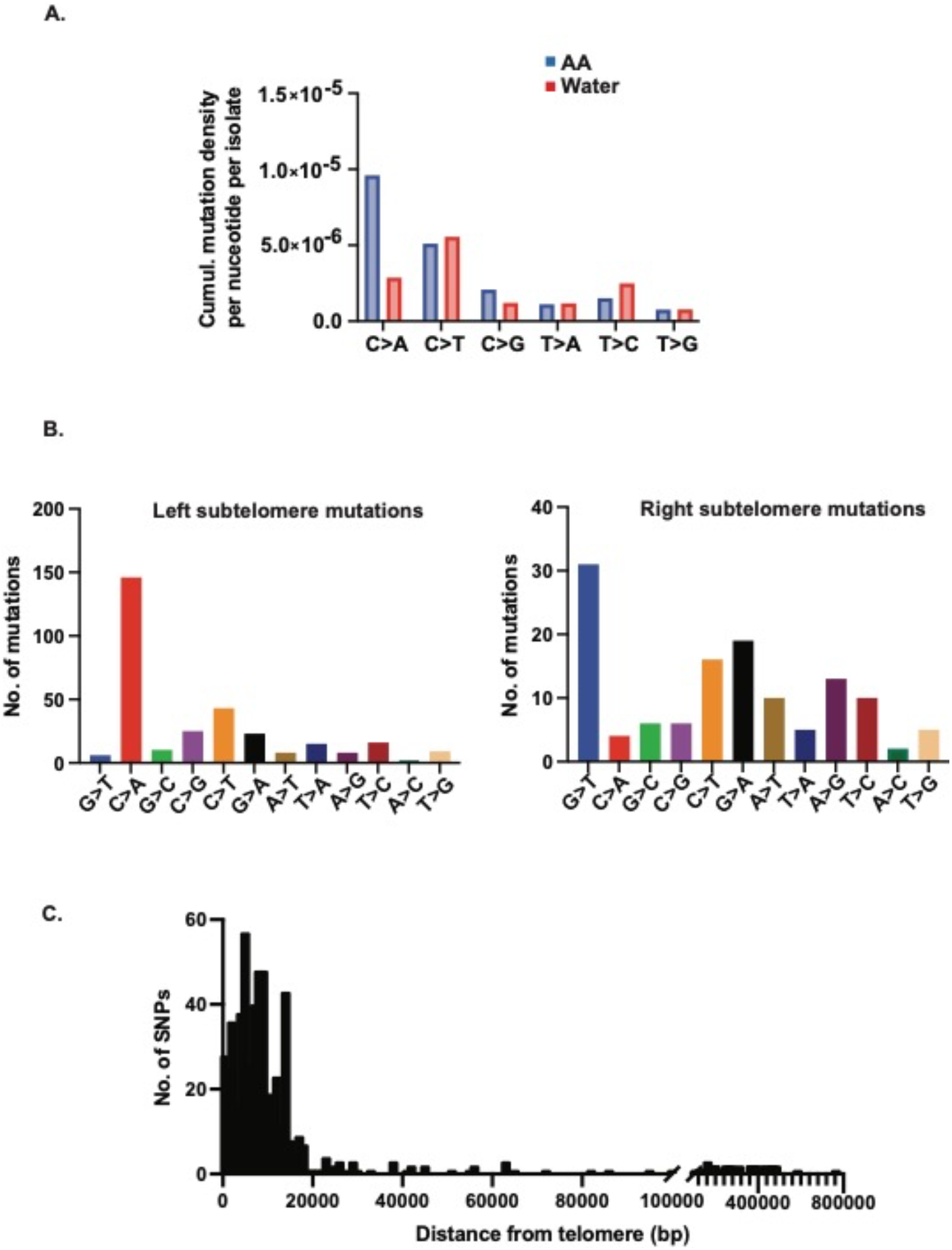
Mutation spectrum of acetaldehyde in yeast. Whole-genome sequenced Can^R^Ade^-^ mutants were aligned to the ySR127 yeast genome (30) using BWA and mutations were called using VarScan A) Cumulative mutation density for each single base substitution (given substitution along with substitution of the complementary base) per isolate with AA or control (water) treatment, calculated as the number of given substitutions per the total number of mutable bases within ssDNA (30 kb from telomeres). B) Frequency of base substitutions near left and right telomeres to assess bias of substitutions. C) Distance of mutations from telomeres to ascertain sub-telomeric vs mid-chromosome distribution. Chromosome coordinates for yeast reference sequence (sacCer3) were obtained from UCSC Table Browser and distances were estimated using BEDtools. Plotting and analysis were performed using Prism (v 9.3.1, GraphPad Software, LLC).

In our assay, 5’→3’ resection from chromosome ends would render the bottom strand of the left arm and top strand of the right chromosomal arm single stranded. Accordingly, we asked if AA prefers mutating a specific base on single stranded DNA. Upon comparing the locations of the observed mutations to the reference strand, we observed that C→A changes were pre-dominant on the left arms of chromosomes (indicating bottom single-strand G lesions) and conversely saw G→T changes on the right chromosome arms (indicating top single-strand G lesions) (Fig 2B). We estimated the distance of mutations from the telomeres and observed that most mutations (453/504, Table S4) clustered within 30 kb of the telomeres (Fig 2C), while ~12% (53/504, Table S4) mutations were found in the mid-chromosomal regions. Further, we noticed that most of the unselected AA-derived mutations (i.e mutations excluding chromosome V) were within 30 kb from telomeres (224/240, Table S4), indicating that AA has a propensity to damage ssDNA regardless of a selection bias. Our results confirm that in our assay system, telomere-proximal single-stranded DNA is the preferred template for AA-induced mutagenesis.

Overall, our analysis reveals a distinct mutation spectrum for AA, whereby guanines are strongly preferred over other bases and the ensuing G→T (C→A) transversions constitute the major mutation type. We did not observe enough indels in our samples for signature analysis (Table S4). Nevertheless, our data agrees with prior *in vitro* studies showing that the primary target of AA induced DNA damage are guanine residues (8,14,46).

### Mutation signature of acetaldehyde in single stranded DNA

We next determined the proportion of mutations falling within all the 96 possible single base substitutions within trinucleotide mutational motifs, consisting of a central base and the flanking −1 and +1 residues (33). When plotted as cumulative pyrimidine changes (i.e., mutated base along with the complementary mutated base) there was a marked enrichment of C→A mutations when the −1 base was a guanine (Fig 3A, Table S5). The contribution of the remaining signature motifs remained at the baseline level, strongly suggesting that the major mutation signature of AA is centered around mutated guanine residues (showing as C→A changes).

**Fig 3:**
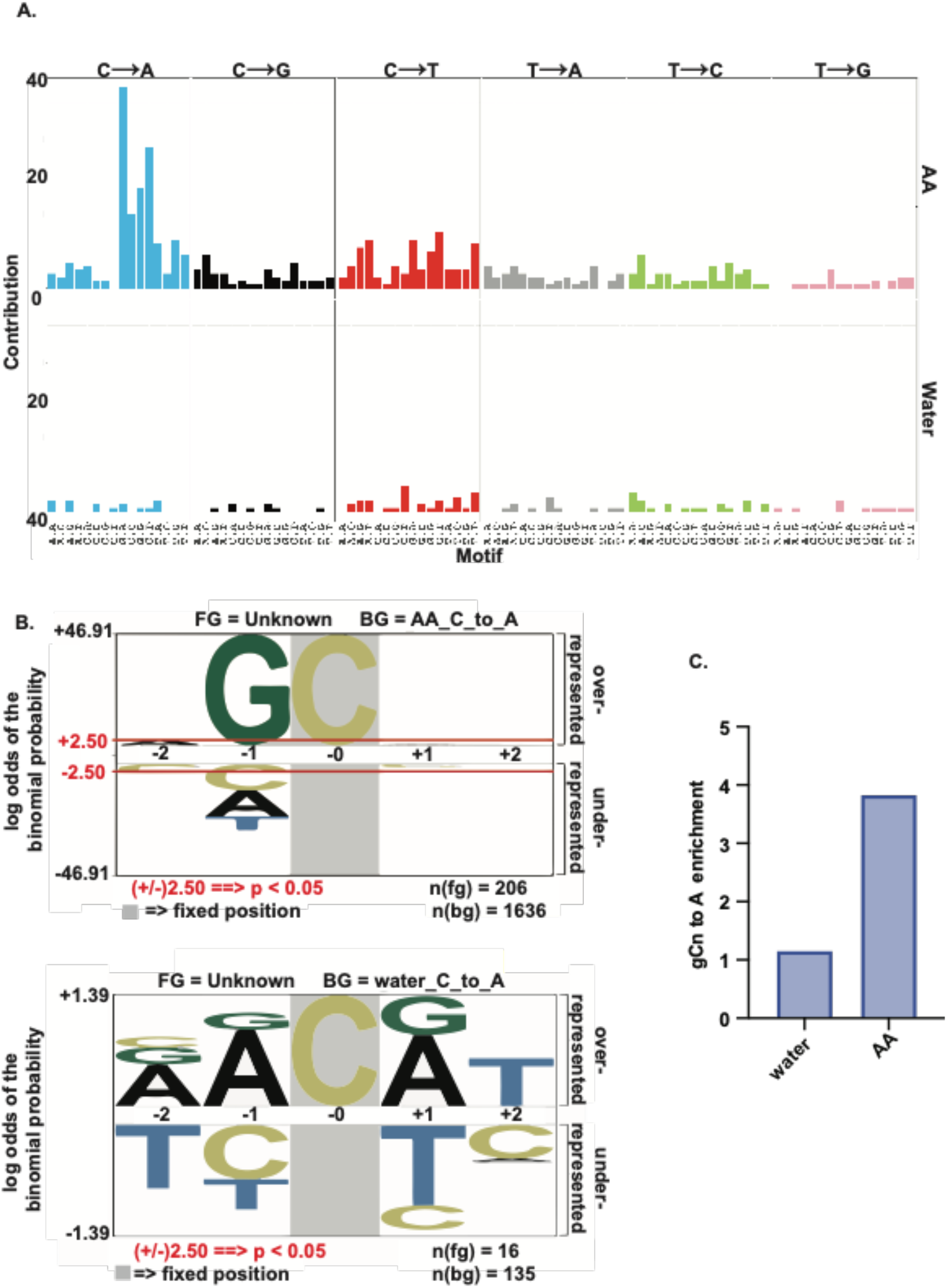
Acetaldehyde mutation signature analysis. A) Contribution of each of the 96 possible base substitutions to the AA mutation signature are depicted. For comparison, mutations from AA (top) and control (water, bottom) samples were analyzed concomitantly. The analysis was performed using the Bioconductor R package SomaticSignatures (33). B) Over-representation of nucleotides in a pentanucleotide context of C→A mutations using pLogo. Cytosine in grey highlight represents the fixed C position and heights of residues in the −2 to +2 positions indicate magnitude of over- or underrepresentation of the indicated residue at the position. N(fg)= foreground mutations i.e total number of C→A substitutions. N(bg)= background mutations i.e number of all other C substitutions across the genome. Red lines in top panel represent over/under-represented residues that are statistically significant. C) Enrichment of the gCn→gAn AA mutation signature in yeast.

To confirm the above observation, we used pLogo to check the proportion of over-and under-represented nucleotides flanking the C→A (or G→T) change. We noticed a strong over-representation for a guanine in the −1 position flanking the mutated cytosine (Fig 3B, Table S6 conversely represented to show mutated base as a pyrimidine), yielding a gCn→gAn (nGc→nTc) signature for AA exposure. In comparison, no statistical enrichment was observed for this signature in the water-treated control samples (Fig.3B). We further used the knowledge-based pipeline we termed TriMS (*T*rinucleotide *M*utation *S*ignature) to determine if this mutation signature is enriched in our samples. TriMS is broadly customizable to any oligonucleotide-centered mutational motif. We calculated the enrichment of the gCn→gAn change versus all C→A changes genome-wide, with an enrichment ≥ 1 and a Benjamini-Hochberg-corrected p-value ≤0.05 deemed statistically significant. In the AA-treated samples, the gCn→gAn signature was highly enriched (3.8, BH-corrected p-value 2.09E-22) compared to control samples (1.15 BH-corrected p-value 0.65829269) (Fig 3C, Table S7).

### Acetaldehyde-derived mutation signature can be found in alcohol-associated human cancers

Alcohol consumption is unequivocally associated with the risk of malignancy for a variety of cancer types, including cancers of the digestive tract, colorectal cancer as well as hepatocellular carcinomas (47). Since *AA* concentrations are likely elevated in alcohol-associated cancers, we sought to detect the AA mutation signature in published cancer datasets. In addition, AA is present in tobacco smoke (48), thus we also analyzed lung cancers for enrichment of the AA mutation signature.

We analyzed >1800 whole-exome sequenced cancers spanning 5 alcohol- or smoking-associated cancer types from ICGC and >500 whole-genome sequenced cancers across 5 alcohol-associated cancer types from the PCAWG database. We identified a significant enrichment of the gCn→gAn (nGc→nTc) signature in various samples amongst both cohorts (Fig. 4A, Table S8). In the whole-exome sequenced lung cancer samples, we detected signature enrichment at a low frequency (5/1001 samples with enrichment >1 for LUAD and LUSC, Fig 4A). In liver cancers, 27 samples across 3 different cancer cohorts (LICA-FR, LIHC, LIRI-JP) had a significant gCn→gAn enrichment (Fig 4A). Within the whole-genome sequenced PCAWG datasets, we were able to identify samples from esophageal carcinoma (ESCA) and head-and-neck cancer (HNSCC) that had a significant enrichment of the gCn→gAn mutation signature ((>1, BH-corrected p-value ≤ 0.05), Fig. 5A, Table S10). Like the ICGC samples, various whole-genome sequenced hepatocellular carcinoma, as well as stomach adenocarcinoma samples demonstrated a statistically significant enrichment of the AA-associated mutation signature gCn→gAn (Fig 5A). We also analyzed >7000 samples spanning 19 additional cancers in the whole-exome sequenced ICGC datasets, and ~1100 samples spanning 11 additional whole-genome sequenced PCAWG cancers. We did not see an enrichment for the AA signature in reproductive, neurological, renal, or hematological cancers (Tables S12, S13).

**Fig 4:**
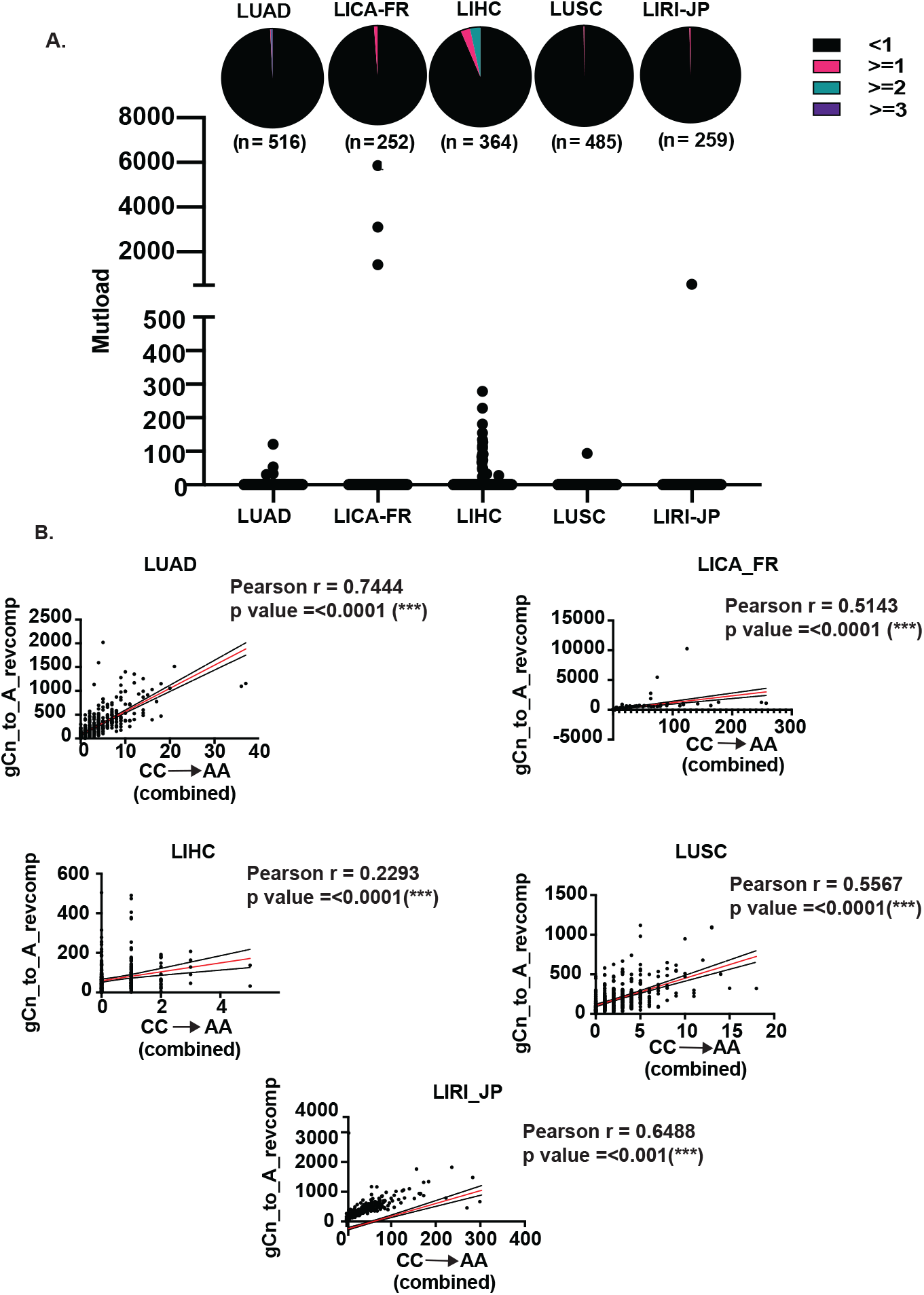
Acetaldehyde mutation spectrum in whole-exome sequenced ICGC cancers associated with smoking/alcohol. A) Scatterplot depicting mutation loads in samples displaying a fold enrichment of the AA mutation signature ≥1 with a Benjamini-Hoechberg corrected p-value of ≤0.05. Samples displaying enrichment are represented as colored sectors in the pie chart situated above the corresponding mutation loads for each cancer dataset. The total number of samples analyzed per cohort is in parentheses under the pie charts. B) Correlation of the CC→AA dinucleotide mutation signature with gCn→gAn trinucleotide mutation signature in ICGC cancers from (A). For each dataset, black dots represent the gCn→gAn mutation loads for each sample, red line is the linear regression and black lines are the 95% confidence intervals. Plotting and analysis were performed using Prism (v 9.3.1, GraphPad Software, LLC) and R Studio (http://www.rstudio.com/).

**Fig 5:**
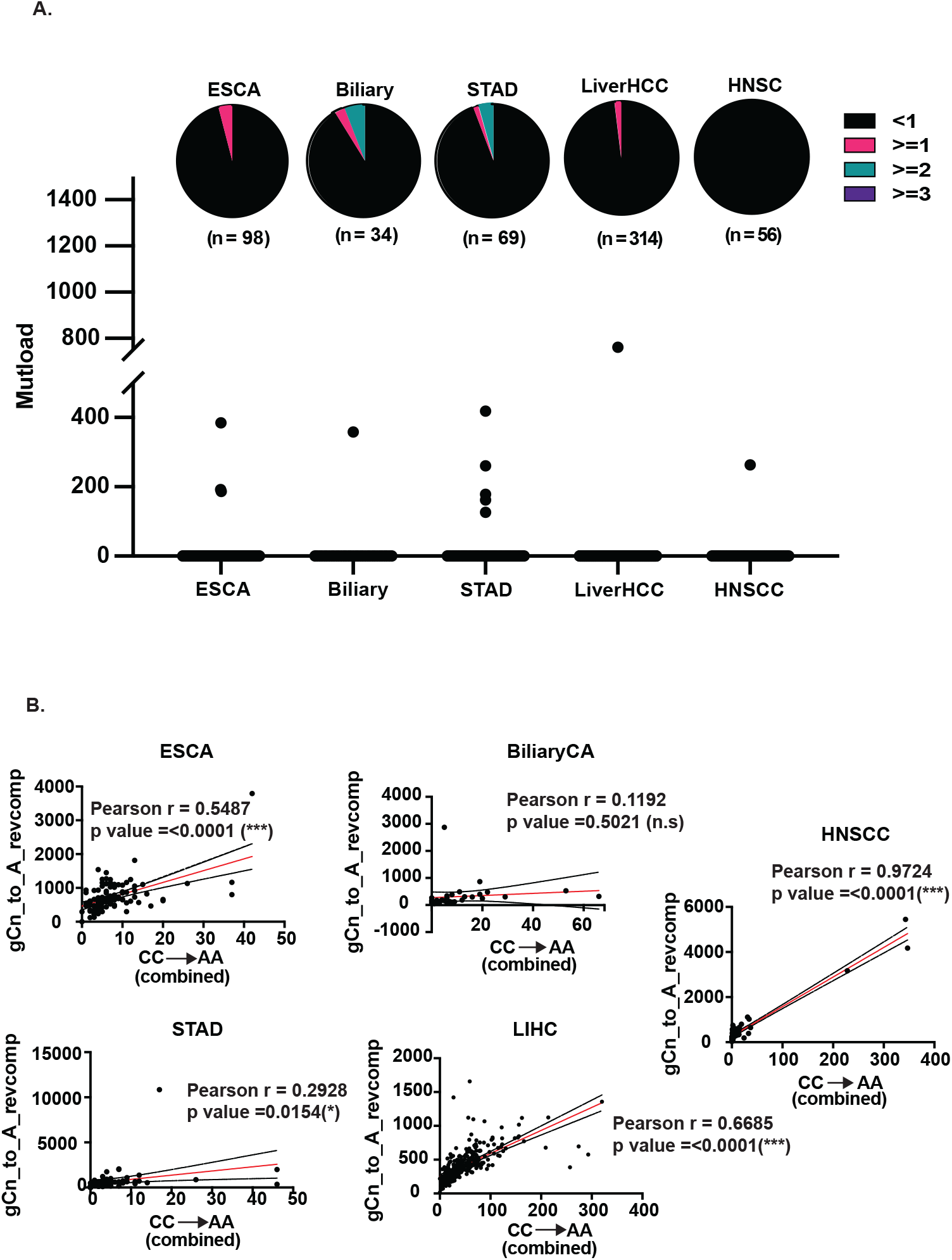
Acetaldehyde mutation spectrum in whole-genome sequenced PCAWG cancers associated with smoking/alcohol. A) Scatterplot depicting mutation loads in samples displaying a fold enrichment of the AA mutation signature ≥1 with a Benjamini-Hoechberg corrected p-value of ≤0.05. Samples displaying enrichment are represented as colored sectors in the pie chart situated above the corresponding mutation loads for each cancer dataset. The total number of samples analyzed per cohort is in parentheses under the pie charts. B) Correlation of the CC→AA dinucleotide mutation signature with gCn→gAn trinucleotide mutation signature in PCAWG cancers from (A). For each dataset, black dots represent the gCn→gAn mutation loads for each sample, red line is the linear regression and black lines are the 95% confidence intervals. Plotting and analysis were performed using Prism (v 9.3.1, GraphPad Software, LLC) and R Studio (http://www.rstudio.com/).

AA exposure is typically associated with an increase in CC→AA (or GG→TT) transversions (14,49). We hypothesized since gCn→gAn mutations are also induced by AA, CC→AA mutation loads should correlate with the gCn→gAn mutations in AA-associated cancer types. We observed that for most of the ICGC and PCAWG cohorts listed above, there was a positive correlation between the cumulative CC→AA double base substitutions and the cumulative gCn→gAn mutation loads in the same samples (Fig 4B, Fig 5B, Tables S9, S11). We surmise that acetaldehyde can produce at least two distinct mutational signatures in alcohol-associated cancers,

Lastly, we looked for clinical correlations between alcohol/smoking and the presence of the AA-associated mutation signature. The absence of comprehensive clinical data for the ICGC samples and most of the PCAWG samples, particularly with regards to alcohol and tobacco history, precluded analysis of correlations between signature enrichment and patient exposure. However, within the PCAWG esophageal cancer (ESCA) dataset, 5 samples that were enriched for the gCn→gAn signature also had associated clinical data with all samples demonstrating significant enrichment belonging to patients with a known history of either tobacco usage or alcohol consumption (Fig S2, Table S14).

### The acetaldehyde mutation signature displays transcriptional strand bias in human cancers

During transcription, the template strand (transcribed strand) remains associated with the newly synthesized RNA molecule, and lesions on this strand are capable of stalling RNA polymerase, which leads to the recruitment of transcription-coupled repair machinery such as TC-NER (50,51). Conversely, the coding strand (non-transcribed strand) is rendered single stranded, which makes it particularly susceptible to transcription-associated mutagenesis (52). Many cancers display a strand asymmetry of mutations, for example those associated with UV exposure, smoking, alkylating agents, or oxidative damage (29,53,54). Exploring such mutational biases can greatly contribute to the overall understanding of the mutational processes that drive carcinogenesis. To this end, we asked if the cancers displaying significant enrichment for the gCn→gAn (nGc→nTc) mutation signature have a transcriptionally biased mutation spectrum. For whole-exome sequenced, liver-associated cancers from ICGC, there was strong bias for nGc→nTc to occur on the non-transcribed strand (Fig 6A, Table S15). Interestingly, we did not observe a statistically significant bias for lung-associated cancers (Fig 6B, Table S15). Similarly in wholegenome sequenced PCAWG cancers datasets, significantly more nGc→nTc mutations were found associated with the non-transcribed strand in liver and stomach-associated cancers, but not in upper respiratory tract-associated ESCA and HNSCC (Fig 6B, Table S15). Our results suggest that elevated transcription in cancer cells would generate abundant ssDNA and likely provide more opportunities for AA-induced damage, especially in liver cancers which are often associated with chronic alcohol exposure.

**Fig 6:**
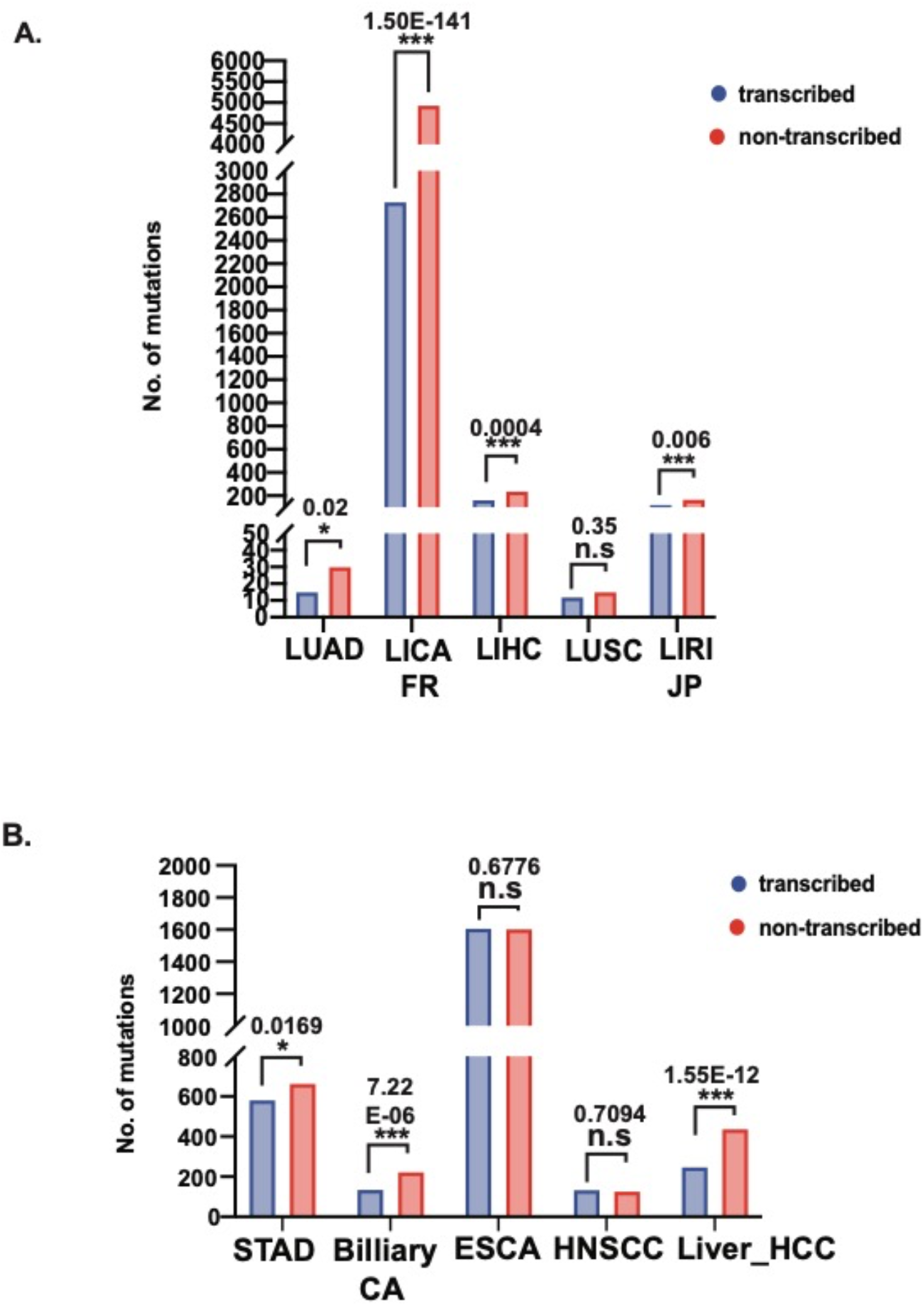
Transcriptional strand bias of the acetaldehyde-associated mutation signature in cancers. Calculations were performed in cancer cohorts displaying a statistically significant fold enrichment of the AA mutation signature. Benjamini-Hoechberg corrected p-values indicate whether the strand bias is statistically significant. A) IAGC cancers, B) PCAWG cancers.

## Discussion

### Acetaldehyde mutagenicity and signature

In the present study, we demonstrated that acetaldehyde (AA) exposure generates strand-biased, guanine centered mutations upon damage in ssDNA in yeast. We showed that AA is highly mutagenic on ssDNA and observe a preponderance of C→A (G→T) single nucleotide polymorphisms. We surmise the observed mutations likely arise from lesion bypass by the Pol ζ polymerase associated with error-prone translesion synthesis. We deciphered a distinct gCn→gAn (nGc→nTc) mutation signature for AA in yeast. Importantly, we were able to detect an enrichment of the AA-associated signature in various alcohol-associated cancer genomes, indicating that the mutation signature identified in yeast is diagnostic of AA-exposure in cancers.

Previously GG→TT changes have been ascribed to AA-induced DNA damage in *in vitro* studies (14,49). Interestingly, we did not see an enrichment in GG→TT changes or other double base substitutions in yeast strains treated with AA. The difference in the signature of AA-induced mutagenesis likely reflects the type of lesions formed in double-stranded plasmid DNA *in vitro* as compared to the lesions in ssDNA *in vivo.* Two molecules of AA can react with guanines to form CrPdG (55). When the open-ring forms of CrPdG react with each other, inter- or intra-strand crosslinks may be formed (14,56,57). Mutagenic bypass of these crosslinks leads to GG→TT changes— double base substitutions classically associated with AA exposure (14,18,22). On the other hand, various studies have demonstrated that AA also forms lesions on single guanine moieties, with N2-ethylidene-dG being the most commonly derived adduct (58), and bypass of such lesions results in single base substitutions at guanine residues (9,10,12). It is possible that AA predominantly forms such mutagenic lesions on single guanine residues in ssDNA in yeast, leading to G→T single base substitutions.

Furthermore, studies in fission yeast have demonstrated that AA exposure leads to the activation of various DNA repair pathways including nucleotide excision repair, base excision repair and homologous recombination (17). As such, it is likely that a majority of the studies aimed at understanding the mutation signatures of AA were unable to detect single base substitutions, as the N2-ethylidene-dG lesion could have been efficiently removed by these DNA repair pathways. Because excision repair pathways cannot function on ssDNA, we are likely able to enrich and reliably detect single base substitutions associated with erroneous bypass of this lesion in our system.

Based on prior reports of a link between ethanol consumption and oxidative stress (59), it is reasonable to assume that AA treatment by itself may also impart oxidative stress. In yeast, induction of oxidative stress has been shown to produce a distinct C→T mutation signature in ssDNA (54). However, in our yeast samples, we observe a much lower number of sub-telomeric C→T mutations compared to C→A changes with AA treatment (Table S4). Although we cannot fully rule out AA-induced oxidative stress contributing to the observed C→A mutagenesis, our data suggests that this most likely is not the primary mechanism of AA-associated mutagenesis.

### Bypass of acetaldehyde-induced DNA damage by TLS in yeast

Mutagenic bypass of DNA lesions requires the activity of translesion polymerases. Studies on alcohol-associated cancers identified mutation signature associated with the TLS polymerase Pol η (60). Additionally, alcohol-induced mutagenesis in budding yeast and AA sensitivity in fission yeast was found to be dependent on the activity of translesion polymerases (17,24). Further, removal of Rev1 in cell-free assays has been shown to impact the mutagenicity of AA-derived interstrand crosslinks (18). In agreement with the above, we observed that TLS was required for AA-induced mutagenesis. Abolishing *REV3* led to a drastic reduction in AA-induced mutation frequency (Fig 1, Table S2), indicating that Polζ is essential for the mutagenic bypass of AA-induced lesions in ssDNA in budding yeast.

We detected an enrichment of the AA-associated mutation signature (gCn→gAn) in several samples from alcohol-associated cancers in the ICGC and PCAWG datasets; however, no significant enrichment for the AA-associated mutation signature was seen in any of the several other whole-genome and -exome sequenced cancer datasets, including reproductive cancers, skin malignancies, neurological cancers as well as urothelial cancers (Table S12, S13). As such, it is reasonable to argue that chronic alcohol exposure leads to higher AA-induced genomic damage, especially in the form of lesions on ssDNA significantly contributes to, carcinogenesis. The observed correlation between the smoking/drinking status and AA signature enrichment for esophageal carcinoma samples further substantiates this argument (Fig S2). However, the lack of comprehensive clinical data for different cancer types prevents better statistical analysis of the correlation between chronic alcohol consumption and AA-induced mutation signature in these cancers. Notably, in cohorts from ICGC and PCAWG cancers associated with alcohol consumption, we observed a remarkable correlation between an increase in CC→AA double base substitutions and elevated gCn→gAn mutations (Figs. 4,5). Our data underlines the specificity of the acetaldehyde-associated mutation signature and suggests that *in vivo*, AA mutagenesis likely occurs in a repair-, and template-dependent manner, with differential lesions on ssDNA vs dsDNA resulting in varying mutation outcomes.

For both PCAWG and ICGC cancers analyzed in our study, we observe a transcriptional strand bias for the AA mutation signature; however, this bias is more pronounced in cancers typically associated with heavy alcohol consumption, mainly liver and/or gastro-intestinal tumors (Fig 6). Surprisingly, we see either no strand bias (LUSC, ESCA, HNSCC) or a small degree of bias (LUAD) in cancers of the upper respiratory tract, even though these tissue types are sites of primary exposure to ethanol. One possibility is that most of consumed ethanol is metabolized in hepatocytes, which ensures a higher probability of exposure to AA in hepatic and surrounding tissues (61,62). Also, oral AA levels are influenced by a gamut of factors including beverage type, tobacco smoking history, oral hygiene, and metabolism via the oral microbiome (63–66). The variability in AA exposure on oral and upper respiratory tissues could alter the genome-wide distribution, accumulation and/or spectra of mutations associated with AA exposure in these tissues. On the other hand, oral and upper-respiratory tract tissues are also exposed to a wide variety of other mutagens including tobacco smoke which can lead to an accumulation of lesions and mutations in guanines leading the characteristic C→A (G→T) changes (22,67). Such overlapping mutations may confound the analysis of the contribution of AA-induced mutations in these samples.

Previously, a T→C mutation signature (Signature E4/SBS Signature 16) was ascribed to alcohol and smoking in esophageal carcinoma (26,68), however the molecular etiology of the signature was unknown. Previous studies have described a smoking associated signature in COSMIC (SBS Signature 4), and tobacco chewing (SBS Signature 29), and reactive oxygen species (ROS) (SBS Signature 18) (22), which have a similar mutation pattern to that we observed for AA, predominantly C→A changes. However, unlike our analysis, there was no clear trinucleotide mutational motif in these studies. As such, these signatures often mutations from other etiologies and are not diagnostic of AA exposure.

## Conclusion

Environmental aldehydes represent a growing class of toxic agents that are linked with an increasing risk of many human ailments, including neurodegenerative disease, cardiopulmonary diseases, and aging. Due to similarities in their physicochemical properties, environmental and endogenous aldehydes can not only act synergistically but also can cross-react to produce amplified genotoxic effects (69). The variability in reported AA-associated mutagenesis from past studies in model systems and in alcohol-associated cancers suggest that the AA mutation spectrum might be governed by multiple factors, including specific genomic contexts, the replication/transcriptional status, DNA repair proficiency, and perhaps epigenetic modifications. Understanding the molecular mechanisms underpinning aldehyde toxicity would go a long way in determining the risks associated with exposure to addictive agents such as alcohol and tobacco smoke and devising appropriate therapeutic strategies.

## Data and code availability

The yeast strains used in the study are available upon request. Raw FASTQ sequence files from wholegenome sequencing of yeast samples have been deposited to the Sequence Read Archives (SRA) database under BioProject ID PRJNA817061. Sequence for the reference yeast genome used in this study (ySR127) is accessible on GenBank (CP011547-CP011563). Source code for TriMS is available on GitHub (https://github.com/nataliesaini11/TriMS).

## Acknowledgements

We would like to thank D.Gordenin, J.Delaney and members of the Saini lab for their critical reading of the manuscript and helpful comments.

## Author contributions

S.V, N.S conceptualized the study and designed experiments, S.V, L.S performed the experiments, P.A.M performed whole-genome sequencing of yeast samples, S.V, N.S and L.P analyzed the data. S.V and N.S wrote the manuscript.

---

**Fig S1:**
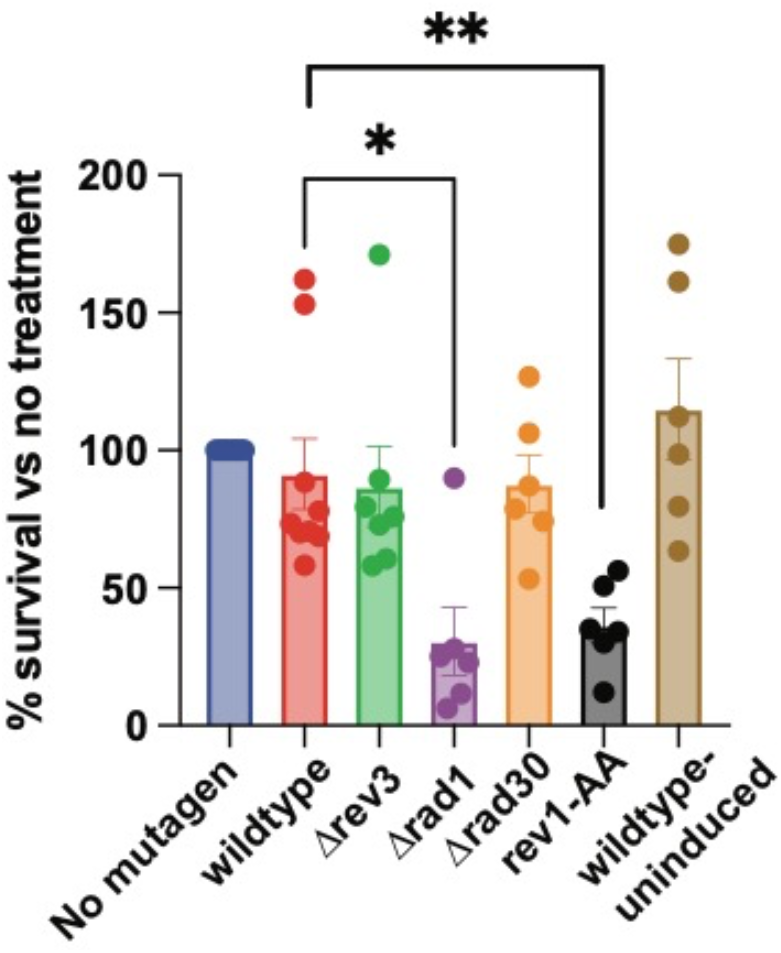
Viability of the strains from Figure 1 with acetaldehyde treatment, shown as percentage of survival compared to the corresponding control treated samples (100%). Data is shown as mean survival with error bars representing standard error of mean. Asterisks indicates statistical significance based on a p-value <0.05 from an unpaired t-test (wildtype vs Δ*rad1*= 0.01, wildtype vs *rev1-AA*= 0.005)

**Fig S2:**
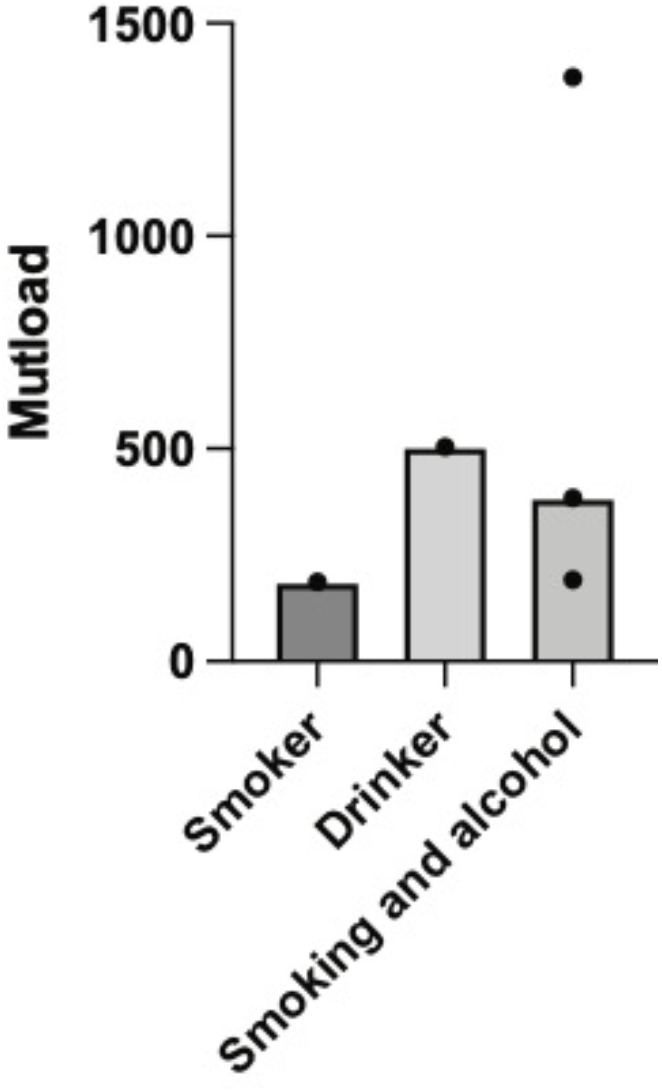
Clinical correlation of acetaldehyde mutation loads with smoking/alcohol status. Whole-genome sequenced esophageal carcinoma samples (ESCA) from PCAWG showing a statistically significant fold enrichment of the AA mutation signature were classified according to smoking/alcohol usage against gCn→gAn mutation loads within these samples. All samples with enrichment of the acetaldehyde signature were either smokers, drinkers, or both.

## Supplementary table legends

Table S1: Yeast strains and oligonucleotide primers used in the study.

Table S2: Pilot data testing acetaldehyde mutagenesis in yeast ssDNA reporter system, along with source data for Fig 1 showing Can^R^Ade^-^ mutation frequencies in mutant yeast strains treated with AA or control.

Table S3: Phenotypic tests for whole-genome sequenced yeast strains treated with acetaldehyde or water.

Table S4: Source data for Fig 2. For samples ± acetaldehyde, single base substitutions, combined unique indels, mutation density per isolate, distribution of mutations in left and right sub-telomeric regions, distance of mutations from telomeres, and mutation spectrum per isolate for water vs AA-treated samples are shown.

Table S5: Source data for Fig 3A showing the number of mutations within all possible 96 trinucleotide contexts.

Table S6: Source data for Fig 3B showing the pentanucleotide sequence contexts for PLogo analysis.

Table S7: Source data for Fig 3C showing analysis of enrichment for the gCn to gAn mutation signature in whole genome sequenced yeast.

Table S8: Source data for Fig 4A showing analysis of enrichment for the gCn to gAn mutation signature in whole exome sequenced ICGC cancers. Only cancers with samples having an enrichment of ≥1 (BH-corrected Fisher’s p-value ≤0.05) are shown.

Table S9: Source data for Fig 4C showing correlation between the AA gCn→gAn signature and cumulative CC→AA (CC→AA + GG→TT) dinucleotide base substitutions in the indicated ICGC cancer cohorts.

Table S10: Source data for Fig 5A showing analysis of enrichment for the gCn to gAn mutation signature in whole genome sequenced PCAWG cancers. Only cancers with samples having an enrichment of ≥1 (BH-corrected Fisher’s p-value ≤0.05) are shown.

Table S11: Source data for Fig 5C showing correlation between the AA gCn→gAn signature and cumulative CC→AA (CC→AA + GG→TT) dinucleotide base substitutions in the indicated PCAWG cancer cohorts.

Table S12: Analysis of enrichment for the gCn to gAn mutation signature in whole exome sequenced ICGC cancers for cancers not displaying an enrichment of ≥1 (BH-corrected Fisher’s p-value ≤0.05).

Table S13: Analysis of enrichment for the gCn to gAn mutation signature in whole genome sequenced PCAWG cancers for cancers not displaying an enrichment of ≥1 (BH-corrected Fisher’s p-value ≤0.05).

Table S14: Source data for Fig S2 showing the tobacco and alcohol exposure clinical data for wholegenome sequenced ESCA (esophageal carcinoma) samples from PCAWG.

Table S15: Source data for Fig 6A and 6B showing the transcriptional strand bias of the gCn-to-gAn acetaldehyde mutation signature in ICGC and PCAWG cancers displaying an enrichment of the AA mutation signature.

## References

1. Mizumoto, A., Ohashi, S., Hirohashi, K., Amanuma, Y., Matsuda, T. and Muto, M. (2017) Molecular Mechanisms of Acetaldehyde-Mediated Carcinogenesis in Squamous Epithelium. Int J Mol Sci, 18.

2. Vasiliou, V. and Nebert, D.W. (2005) Analysis and update of the human aldehyde dehydrogenase (ALDH) gene family. Hum Genomics, 2, 138–143.

3. Shortall, K., Djeghader, A., Magner, E. and Soulimane, T. (2021) Insights into Aldehyde Dehydrogenase Enzymes: A Structural Perspective. Front Mol Biosci, 8, 659550.

4. Heymann, H.M., Gardner, A.M. and Gross, E.R. (2018) Aldehyde-Induced DNA and Protein Adducts as Biomarker Tools for Alcohol Use Disorder. Trends Mol Med, 24, 144–155.

5. Tsuruta, H., Sonohara, Y., Tohashi, K., Aoki Shioi, N., Iwai, S. and Kuraoka, I. (2020) Effects of acetaldehyde-induced DNA lesions on DNA metabolism. Genes Environ, 42, 2.

6. Humans, I.W.G.o.t.E.o.C.R.t. (2012) Personal habits and indoor combustions. Volume 100 E. A review of human carcinogens. IARC Monogr Eval Carcinog Risks Hum, 100, 1–538.

7. Brooks, P.J. and Zakhari, S. (2014) Acetaldehyde and the genome: beyond nuclear DNA adducts and carcinogenesis. Environ Mol Mutagen, 55, 77–91.

8. Wang, M., McIntee, E.J., Cheng, G., Shi, Y., Villalta, P.W. and Hecht, S.S. (2000) Identification of DNA adducts of acetaldehyde. Chem Res Toxicol, 13, 1149–1157.

9. Matter, B., Guza, R., Zhao, J., Li, Z.Z., Jones, R. and Tretyakova, N. (2007) Sequence distribution of acetaldehyde-derived N2-ethyl-dG adducts along duplex DNA. Chem Res Toxicol, 20, 1379–1387.

10. Terashima, I., Matsuda, T., Fang, T.W., Suzuki, N., Kobayashi, J., Kohda, K. and Shibutani, S. (2001) Miscoding potential of the N2-ethyl-2’-deoxyguanosine DNA adduct by the exonuclease-free Klenow fragment of Escherichia coli DNA polymerase I. Biochemistry, 40, 4106–4114.

11. Perrino, F.W., Blans, P., Harvey, S., Gelhaus, S.L., McGrath, C., Akman, S.A., Jenkins, G.S., LaCourse, W.R. and Fishbein, J.C. (2003) The N2-ethylguanine and the O6-ethyl-and O6-methylguanine lesions in DNA: contrasting responses from the “bypass” DNA polymerase eta and the replicative DNA polymerase alpha. Chem Res Toxicol, 16, 1616–1623.

12. Upton, D.C., Wang, X., Blans, P., Perrino, F.W., Fishbein, J.C. and Akman, S.A. (2006) Replication of N2-ethyldeoxyguanosine DNA adducts in the human embryonic kidney cell line 293. Chem Res Toxicol, 19, 960–967.

13. Cheng, T.F., Hu, X., Gnatt, A. and Brooks, P.J. (2008) Differential blocking effects of the acetaldehyde-derived DNA lesion N2-ethyl-2’-deoxyguanosine on transcription by multisubunit and single subunit RNA polymerases. J Biol Chem, 283, 27820–27828.

14. Matsuda, T., Kawanishi, M., Yagi, T., Matsui, S. and Takebe, H. (1998) Specific tandem GG to TT base substitutions induced by acetaldehyde are due to intra-strand crosslinks between adjacent guanine bases. Nucleic Acids Res, 26, 1769–1774.

15. Mechilli, M., Schinoppi, A., Kobos, K., Natarajan, A.T. and Palitti, F. (2008) DNA repair deficiency and acetaldehyde-induced chromosomal alterations in CHO cells. Mutagenesis, 23, 51–56.

16. Brendel, M., Marisco, G., Ganda, I., Wolter, R. and Pungartnik, C. (2010) DNA repair mutant pso2 of Saccharomyces cerevisiae is sensitive to intracellular acetaldehyde accumulated by disulfiram-mediated inhibition of acetaldehyde dehydrogenase. Genet Mol Res, 9, 48–57.

17. Noguchi, C., Grothusen, G., Anandarajan, V., Martinez-Lage Garcia, M., Terlecky, D., Corzo, K., Tanaka, K., Nakagawa, H. and Noguchi, E. (2017) Genetic controls of DNA damage avoidance in response to acetaldehyde in fission yeast. Cell Cycle, 16, 45–58.

18. Hodskinson, M.R., Bolner, A., Sato, K., Kamimae-Lanning, A.N., Rooijers, K., Witte, M., Mahesh, M., Silhan, J., Petek, M., Williams, D.M. et al. (2020) Alcohol-derived DNA crosslinks are repaired by two distinct mechanisms. Nature, 579, 603–608.

19. Tacconi, E.M., Lai, X., Folio, C., Porru, M., Zonderland, G., Badie, S., Michl, J., Sechi, I., Rogier, M., Matia Garcia, V. et al. (2017) BRCA1 and BRCA2 tumor suppressors protect against endogenous acetaldehyde toxicity. EMBO Mol Med, 9, 1398–1414.

20. Nik-Zainal, S., Alexandrov, L.B., Wedge, D.C., Van Loo, P., Greenman, C.D., Raine, K., Jones, D., Hinton, J., Marshall, J., Stebbings, L.A. et al. (2012) Mutational processes molding the genomes of 21 breast cancers. Cell, 149, 979–993.

21. Alexandrov, L.B., Nik-Zainal, S., Wedge, D.C., Aparicio, S.A., Behjati, S., Biankin, A.V., Bignell, G.R., Bolli, N., Borg, A., Borresen-Dale, A.L. et al. (2013) Signatures of mutational processes in human cancer. Nature, 500, 415–421.

22. Alexandrov, L.B., Kim, J., Haradhvala, N.J., Huang, M.N., Tian Ng, A.W., Wu, Y., Boot, A., Covington, K.R., Gordenin, D.A., Bergstrom, E.N. et al. (2020) The repertoire of mutational signatures in human cancer. Nature, 578, 94–101.

23. Consortium, I.T.P.-C.A.o.W.G. (2020) Pan-cancer analysis of whole genomes. Nature, 578, 82–93.

24. Voordeckers, K., Colding, C., Grasso, L., Pardo, B., Hoes, L., Kominek, J., Gielens, K., Dekoster, K., Gordon, J., Van der Zande, E. et al. (2020) Ethanol exposure increases mutation rate through error-prone polymerases. Nat Commun, 11, 3664.

25. Kucab, J.E., Zou, X., Morganella, S., Joel, M., Nanda, A.S., Nagy, E., Gomez, C., Degasperi, A., Harris, R., Jackson, S.P. et al. (2019) A Compendium of Mutational Signatures of Environmental Agents. Cell, 177, 821–836 e816.

26. Chang, J., Tan, W., Ling, Z., Xi, R., Shao, M., Chen, M., Luo, Y., Zhao, Y., Liu, Y., Huang, X. et al. (2017) Genomic analysis of oesophageal squamous-cell carcinoma identifies alcohol drinking-related mutation signature and genomic alterations. Nat Commun, 8, 15290.

27. International Cancer Genome, C., Hudson, T.J., Anderson, W., Artez, A., Barker, A.D., Bell, C., Bernabe, R.R., Bhan, M.K., Calvo, F., Eerola, I. et al. (2010) International network of cancer genome projects. Nature, 464, 993–998.

28. Chan, K., Sterling, J.F., Roberts, S.A., Bhagwat, A.S., Resnick, M.A. and Gordenin, D.A. (2012) Base damage within single-strand DNA underlies in vivo hypermutability induced by a ubiquitous environmental agent. PLoS Genet, 8, e1003149.

29. Saini, N., Sterling, J.F., Sakofsky, C.J., Giacobone, C.K., Klimczak, L.J., Burkholder, A.B., Malc, E.P., Mieczkowski, P.A. and Gordenin, D.A. (2020) Mutation signatures specific to DNA alkylating agents in yeast and cancers. Nucleic Acids Res, 48, 3692–3707.

30. Chan, K., Roberts, S.A., Klimczak, L.J., Sterling, J.F., Saini, N., Malc, E.P., Kim, J., Kwiatkowski, D.J., Fargo, D.C., Mieczkowski, P.A. et al. (2015) An APOBEC3A hypermutation signature is distinguishable from the signature of background mutagenesis by APOBEC3B in human cancers. Nat Genet, 47, 1067–1072.

31. Li, H. and Durbin, R. (2009) Fast and accurate short read alignment with Burrows-Wheeler transform. Bioinformatics, 25, 1754–1760.

32. Koboldt, D.C., Chen, K., Wylie, T., Larson, D.E., McLellan, M.D., Mardis, E.R., Weinstock, G.M., Wilson, R.K. and Ding, L. (2009) VarScan: variant detection in massively parallel sequencing of individual and pooled samples. Bioinformatics, 25, 2283–2285.

33. Gehring, J.S., Fischer, B., Lawrence, M. and Huber, W. (2015) SomaticSignatures: inferring mutational signatures from single-nucleotide variants. Bioinformatics, 31, 3673–3675.

34. Quinlan, A.R. and Hall, I.M. (2010) BEDTools: a flexible suite of utilities for comparing genomic features. Bioinformatics, 26, 841–842.

35. Karolchik, D., Hinrichs, A.S., Furey, T.S., Roskin, K.M., Sugnet, C.W., Haussler, D. and Kent, W.J. (2004) The UCSC Table Browser data retrieval tool. Nucleic Acids Res, 32, D493–496.

36. Nugent, C.I., Hughes, T.R., Lue, N.F. and Lundblad, V. (1996) Cdc13p: a single-strand telomeric DNA-binding protein with a dual role in yeast telomere maintenance. Science, 274, 249–252.

37. Saini, N., Zhang, Y., Nishida, Y., Sheng, Z., Choudhury, S., Mieczkowski, P. and Lobachev, K.S. (2013) Fragile DNA motifs trigger mutagenesis at distant chromosomal loci in saccharomyces cerevisiae. PLoS Genet, 9, e1003551.

38. Lee, S.E., Moore, J.K., Holmes, A., Umezu, K., Kolodner, R.D. and Haber, J.E. (1998) Saccharomyces Ku70, mre11/rad50 and RPA proteins regulate adaptation to G2/M arrest after DNA damage. Cell, 94, 399–409.

39. Garvik, B., Carson, M. and Hartwell, L. (1995) Single-stranded DNA arising at telomeres in cdc13 mutants may constitute a specific signal for the RAD9 checkpoint. Mol Cell Biol, 15, 6128–6138.

40. Roberts, S.A., Sterling, J., Thompson, C., Harris, S., Mav, D., Shah, R., Klimczak, L.J., Kryukov, G.V., Malc, E., Mieczkowski, P.A. et al. (2012) Clustered mutations in yeast and in human cancers can arise from damaged long single-strand DNA regions. Mol Cell, 46, 424–435.

41. Yang, Y., Gordenin, D.A. and Resnick, M.A. (2010) A single-strand specific lesion drives MMS-induced hyper-mutability at a double-strand break in yeast. DNA Repair (Amst), 9, 914–921.

42. Marteijn, J.A., Lans, H., Vermeulen, W. and Hoeijmakers, J.H. (2014) Understanding nucleotide excision repair and its roles in cancer and ageing. Nat Rev Mol Cell Biol, 15, 465–481.

43. Haracska, L., Prakash, S. and Prakash, L. (2002) Yeast Rev1 protein is a G template-specific DNA polymerase. J Biol Chem, 277, 15546–15551.

44. Haracska, L., Unk, I., Johnson, R.E., Johansson, E., Burgers, P.M., Prakash, S. and Prakash, L. (2001) Roles of yeast DNA polymerases delta and zeta and of Rev1 in the bypass of abasic sites. Genes Dev, 15, 945–954.

45. Nelson, J.R., Gibbs, P.E., Nowicka, A.M., Hinkle, D.C. and Lawrence, C.W. (2000) Evidence for a second function for Saccharomyces cerevisiae Rev1p. Mol Microbiol, 37, 549–554.

46. Stein, S., Lao, Y., Yang, I.Y., Hecht, S.S. and Moriya, M. (2006) Genotoxicity of acetaldehyde-and crotonaldehyde-induced 1,N2-propanodeoxyguanosine DNA adducts in human cells. Mutat Res, 608, 1–7.

47. Seitz, H.K. and Becker, P. (2007) Alcohol metabolism and cancer risk. Alcohol Res Health, 30, 38–41, 44-37.

48. Seeman, J.I., Dixon, M. and Haussmann, H.J. (2002) Acetaldehyde in mainstream tobacco smoke: formation and occurrence in smoke and bioavailability in the smoker. Chem Res Toxicol, 15, 1331–1350.

49. Sonohara, Y., Yamamoto, J., Tohashi, K., Takatsuka, R., Matsuda, T., Iwai, S. and Kuraoka, I. (2019) Acetaldehyde forms covalent GG intrastrand crosslinks in DNA. Sci Rep, 9, 660.

50. Hanawalt, P.C. and Spivak, G. (2008) Transcription-coupled DNA repair: two decades of progress and surprises. Nat Rev Mol Cell Biol, 9, 958–970.

51. Jiang, G. and Sancar, A. (2006) Recruitment of DNA damage checkpoint proteins to damage in transcribed and nontranscribed sequences. Mol Cell Biol, 26, 39–49.

52. Jinks-Robertson, S. and Bhagwat, A.S. (2014) Transcription-associated mutagenesis. Annu Rev Genet, 48, 341–359.

53. Haradhvala, N.J., Polak, P., Stojanov, P., Covington, K.R., Shinbrot, E., Hess, J.M., Rheinbay, E., Kim, J., Maruvka, Y.E., Braunstein, L.Z. et al. (2016) Mutational Strand Asymmetries in Cancer Genomes Reveal Mechanisms of DNA Damage and Repair. Cell, 164, 538–549.

54. Degtyareva, N.P., Saini, N., Sterling, J.F., Placentra, V.C., Klimczak, L.J., Gordenin, D.A. and Doetsch, P.W. (2019) Mutational signatures of redox stress in yeast single-strand DNA and of aging in human mitochondrial DNA share a common feature. PLoS Biol, 17, e3000263.

55. Garcia, C.C., Angeli, J.P., Freitas, F.P., Gomes, O.F., de Oliveira, T.F., Loureiro, A.P., Di Mascio, P. and Medeiros, M.H. (2011) [13C2]-Acetaldehyde promotes unequivocal formation of 1,N2-propano-2’-deoxyguanosine in human cells. J Am Chem Soc, 133, 9140–9143.

56. Brooks, P.J. and Theruvathu, J.A. (2005) DNA adducts from acetaldehyde: implications for alcohol-related carcinogenesis. Alcohol, 35, 187–193.

57. Cho, Y.J., Wang, H., Kozekov, I.D., Kurtz, A.J., Jacob, J., Voehler, M., Smith, J., Harris, T.M., Lloyd, R.S., Rizzo, C.J. et al. (2006) Stereospecific formation of interstrand carbinolamine DNA cross-links by crotonaldehyde-and acetaldehyde-derived alpha-CH3-gamma-OH-1,N2-propano-2’-deoxyguanosine adducts in the 5’-CpG-3’ sequence. Chem Res Toxicol, 19, 195–208.

58. Matsuda, T., Matsumoto, A., Uchida, M., Kanaly, R.A., Misaki, K., Shibutani, S., Kawamoto, T., Kitagawa, K., Nakayama, K.I., Tomokuni, K. et al. (2007) Increased formation of hepatic N2-ethylidene-2’-deoxyguanosine DNA adducts in aldehyde dehydrogenase 2-knockout mice treated with ethanol. Carcinogenesis, 28, 2363–2366.

59. Comporti, M., Signorini, C., Leoncini, S., Gardi, C., Ciccoli, L., Giardini, A., Vecchio, D. and Arezzini, B. (2010) Ethanol-induced oxidative stress: basic knowledge. Genes Nutr, 5, 101–109.

60. Supek, F. and Lehner, B. (2017) Clustered Mutation Signatures Reveal that Error-Prone DNA Repair Targets Mutations to Active Genes. Cell, 170, 534–547 e523.

61. Paton, A. (2005) Alcohol in the body. BMJ, 330, 85–87.

62. Lieber, C.S. (1997) Ethanol metabolism, cirrhosis and alcoholism. Clin Chim Acta, 257, 59–84.

63. Homann, N., Tillonen, J., Meurman, J.H., Rintamaki, H., Lindqvist, C., Rautio, M., Jousimies-Somer, H. and Salaspuro, M. (2000) Increased salivary acetaldehyde levels in heavy drinkers and smokers: a microbiological approach to oral cavity cancer. Carcinogenesis, 21, 663–668.

64. Stornetta, A., Guidolin, V. and Balbo, S. (2018) Alcohol-Derived Acetaldehyde Exposure in the Oral Cavity. Cancers (Basel), 10.

65. Lachenmeier, D.W., Kanteres, F. and Rehm, J. (2009) Carcinogenicity of acetaldehyde in alcoholic beverages: risk assessment outside ethanol metabolism. Addiction, 104, 533–550.

66. Salaspuro, M.P. (2003) Acetaldehyde, microbes, and cancer of the digestive tract. Crit Rev Clin Lab Sci, 40, 183–208.

67. Nik-Zainal, S., Kucab, J.E., Morganella, S., Glodzik, D., Alexandrov, L.B., Arlt, V.M., Weninger, A., Hollstein, M., Stratton, M.R. and Phillips, D.H. (2015) The genome as a record of environmental exposure. Mutagenesis, 30, 763–770.

68. Li, X.C., Wang, M.Y., Yang, M., Dai, H.J., Zhang, B.F., Wang, W., Chu, X.L., Wang, X., Zheng, H., Niu, R.F. et al. (2018) A mutational signature associated with alcohol consumption and prognostically significantly mutated driver genes in esophageal squamous cell carcinoma. Ann Oncol, 29, 938–944.

69. LoPachin, R.M. and Gavin, T. (2014) Molecular mechanisms of aldehyde toxicity: a chemical perspective. Chem Res Toxicol, 27, 1081–1091.

